# Modulated Zika virus NS1 conjugate offers advantages for accurate detection of Zika virus specific antibody in double antigen binding and Ig capture enzyme immunoassays

**DOI:** 10.1101/603811

**Authors:** Richard S Tedder, Steve Dicks, Samreen Ijaz, Nathalia Caroline Santiago de Souza, Anderson Vincente de Paula, Flavia Levy, Raquel Medialdea-Carrera, José Eduardo Levi, Claudio S Pannuti, Patrícia Carvalho de Sequeira, David W G Brown, Ines Ushiro Lumb

## Abstract

The accurate diagnosis and seroprevalence investigations of Zika virus (ZKV) infections remain complex due to cross reactivity with other flaviviruses. Two assay formats, both using labelled Zika virus NS1 antigen as a revealing agent (a double antigen binding assay, DABA, and an immunoglobulin Ig capture assay, IgG capture) were initially developed and compared with the indirect EuroimmunZ assay for the detection of anti-Zika antibody. Of 147 pre-Zika period serum samples, 39 (27%) were reactive in the EuroimmunZ or the DABA assays, 28 sera concordantly so. Such false reactivity was influenced by the serotype of Dengue virus (DV) to which individuals had been exposed to. Thus, of sera from patients undergoing secondary Dengue virus infection of known serotype, 91%, 45% and 28% of Dengue virus serotype 2, 3 and 4 respectively were reactive in one or more of the three assays. A novel method of quenching false sero-reactivity was therefore developed for the DABA and IgG capture assays. Initial addition of a single homologous Dengue virus serotype 3 NS1Ag quench significantly ablated false reactivities in the pre-Zika period sera. An equipotent quadrivalent quench comprising homologous Dengue virus serotypes 1 to 4 NS1Ag was shown to be optimum yet retained sensitivity for the detection of specific anti-Zika antibody. Comparing DABA and IgG capture assays using quenched and unquenched conjugates in comparison with EuroimmunZ early in the course of PCR-confirmed infection indicated that a significant component of the apparent early anti-ZIKA antibody response is likely to be due to a Zika virus-driven anamnestic anti-Dengue virus response. The increased specificity provided by homologous antigen quenching is likely to provide a significant improvement in sero-diagnostics and to be of clinical value.

## Introduction

Zika virus (ZKV), first described in a nonhuman primate in 1947, is named after a forested area in Uganda [1]. Its ability to infect humans was first demonstrated in 1952 as reviewed by Kindhauser and colleagues [2] with antibody to ZKV detected both by neutralisation and by animal protection experiments [3]. Zika virus is one of a number of closely related Flaviviruses, previously termed Arboviruses, principally transmitted through the bite of an infected *Aedes* mosquito, most commonly *Aedes aegypti* but which may also be transmitted directly between humans through sexual contact or vertically, from mother to the unborn baby. Although possible, no cases of transmission through transplantation of cells, tissues or organs have been reported to date; probable transmission through blood products have been reported [4–6].

Similar to other related flaviruses, ZKV infection is believed to be asymptomatic in up to 80% of cases, with a minority manifesting often mild symptomatology [7]. The clinical illness is of short duration and detectable viraemia seldom exceeds seven days from the illness onset [8]. The infection may persist for longer in a number of sanctuary sites [9] including the male genital tract [4,10].

It is likely that ZKV may have caused outbreaks of disease but that these may well have been unrecognised because the clinical illness is similar to that caused by other known pathogens including Dengue (DV) and Chikungunya viruses. Where ZKV differs is that its neuro-tropism and ability to infect the unborn child transplacentally leads to serious neurological birth defects [11]. The significant morbidity caused by Congenital Zika Syndrome, the societal impact and costs of this in both Polynesia and more recently in countries in South and Central America have added to the requirement for both sensitive and specific tests for the detection of antibody to ZKV (anti-ZKV). However, extensive serological cross-reactivity between flaviviruses and their co-circulation has rendered this difficult to achieve using the conventional indirect immunoassay format.

In collaboration with Corti and his colleagues we previously investigated a novel competitive immunoassay for the detection of anti-ZKV [12]. Whilst it conferred considerably improved accuracy in relation to the limited assay options available, we have further investigated two other test formats for the detection of anti-ZKV including reverse capture (IgG capture) and double antigen binding assay (DABA) principles. Both use a common revealing conjugate of enzyme labelled ZKV NS1 antigen, which together with the addition of un-labelled competitor homologue DV NS1 proteins provides a unique and novel approach to the specific quenching of false reactions due principally to coexisting antibody to other related flaviviruses. Here we describe the performance of these assays and the novel approach to quenching non-specific reactivity in the face of antibody to Dengue viruses as this was the closely-related flavivirus of most relevance in the South American cohorts used in this study, nevertheless recognising that in a global context, other related viruses may induce false serological reactivity. We compared the performance of these two assays with the widely used Euroimmune ZKV antibody Elisa (EuroimmunZ) for the detection of antibody to ZKV in the presence of antibody to Dengue virus (anti-DV) of various serotypes.

## Materials and Methods

### Patient samples

#### Known Dengue types seropositive samples

Sera were available in São Paulo from 134 patients previously infected with DV of PCR confirmed serotype; most samples were taken before 2014, when ZKV is believed to have been introduced into the Americas, including 28 patients with Dengue virus serotype 1 (DV1) primary infection sampled in 2015; 32 patients with Dengue virus serotype 2 (DV2) secondary infection sampled in 2010; 34 patients with Dengue virus serotype 3 (DV3) proven or presumed secondary infection sampled in 2002 and 2005; 40 patients with proven or presumed Dengue virus serotype 4 (DV4) secondary infection sampled in 2013.

#### Pre-Zika period samples

One hundred and forty-seven sera drawn prior to the recognized introduction of ZKV into Brazil were available for testing. One hundred and eighteen were available in Rio de Janeiro from outpatient clinic attenders at Fiocruz during 2014 (outpatient panel) and 29 samples were available in São Paulo from blood donors taken in 2013; 23 of these samples were positive for DV IgG.

#### Zika samples

One hundred and twenty-six sera taken from patients between 1 day and > 1 year after the onset of arboviral-like symptoms of PCR confirmed Zika infection (95 from São Paulo and 31 from Rio de Janeiro) were available for testing.

#### Ethics

In Rio, the procedures applied in this study were performed in accordance with the ethical standards of the Instituto Nacional de Infectologia Evandro Chagas. The study protocol was approved by its Comitê de Ética em Pesquisa (reference CAAE 0026.0.009.000–07). In São Paulo, procedures conformed with terms agreed by the Institutional Review Board from Hospital das Clínicas, University of São Paulo (CAPPesq-Research Projects Ethics Committee). Specimens were anonymised to ensure patient confidentiality.

### Flavivirus antigens

Recombinant ZKV NS1 (rZKVNS1Ag) antigen expressed by baculovirus in insect cells (MyBioSource Inc, San Diego, USA) was used both to coat the solid-phase where appropriate and, when directly conjugated with horseradish peroxidase, to provide a revealing agent for captured antibodies.

Recombinant DV NS1 antigens (rDVNS1Ag) from the four serotypes (DV 1-4 inclusive) expressed in mammalian cells (The Native Antigen Company, Kidlington, Oxfordshire, OX5 1LH, UK) were used in molar excess as components of the conjugate diluent.

### Assay formats

Indirect immunoassay. The EuroimmunZ kit for the detection of IgG anti-ZKV antibody (EUROIMMUN AG, Luebeck, Germany) was used in accordance with the manufacturer’s instructions and defined kit cut-off.

Capture assay. Solid-phase wells (NUNC Immunomodule, U8 Maxisorp wells) were coated with 100μl volumes of Affinipure rabbit anti-γ (Jackson ImmunoResearch, Ely, Cambridgeshire UK) at 5μg/ml in MicroImmune Coating Buffer for ELISA with preservative; (ClinTech, Guildford, UK). Coating was overnight at 2-8°C, followed by 3 hours at 35-37°C. Wells were then washed with PBS Tween 20 and quenched with MicroImmune Blocking Solution (ClinTech, Guildford, UK) for 3-4 hours at 37°C. Wells were aspirated and stored dry at 4°C in sealed pouches with desiccant until use. Prior to testing, serum samples were diluted to 1 in 200 in Transport Medium (TM: Phosphate buffered saline supplemented with Amphotericin B 0.5ug/ml, Gentamicin 0.25mg/ml, 10% v/v heat inactivated foetal calf serum, Tween 20 0.05% v/v, Red Dye 0.05% v/v). One hundred microlitres of diluted serum were added to the wells, incubated for 60 ±2 minutes at 37°C prior to washing and the addition of the conjugate. After a further incubation for 30 ±2 minutes at 37°C the solid-phase was again washed and 100 μl of substrate added, incubated for 30 ±2 minutes at 37°C, the reaction then stopped and measured at 450/630nm. Full details are provided as information for use (IFU) leaflets in supplementary information.

Double antigen binding assay (DABA). Solid-phase wells (NUNC Immunomodule, U8 Maxisorp wells) were coated with 100μl volumes of rZKVNS1Ag at 2μg/ml in MicroImmune Coating Buffer, overnight at 2-8°C, followed by 3 hours at 35-37°C. Wells were then washed with PBS Tween 20 and quenched with MicroImmune Blocking Solution for 3-4 hours at 37°C. Wells were aspirated and stored dry at 4°C in sealed pouches with desiccant until use. Prior to testing 70 μl of sample diluent (MicroImmune Sample Diluent; ClinTech, Guildford, UK) were added to each well. Thirty microlitres of control or test sera were added singly to each well and incubated for 60 ± 2 minutes at 37°C prior to washing and addition of conjugate. After a further incubation for 120 ±5 minutes at 37°C the solid phase was again washed and 100 μl of substrate added, incubated for 30 ±2 minutes at 37°C, the reaction then stopped and measured at 450/630nm. Full details are provided as information for use (IFU) leaflets in supplementary information.

Data from three-way comparisons between EuroimmunZ, the IgG capture and the DABA data generated by individual sera are displayed as Venn diagrams. Two way comparisons are displayed as X by Y plots. Exploration of the efficacy of conjugate performance is also displayed as X by Y plots, comparing conjugate with (blocked) and without (unblocked) the addition of DV NS1 antigens to the conjugate diluent.

### Conjugation of rZKVNS1Ag

One hundred microlitres of recombinant ZIKV NS1 antigen (ensuring at an optimal protein concentration range of 0.5-5.0mg/ml) were coupled to 100μg of lyophilized HRPO mix using the LYNX Rapid HPR Conjugation kit (Bio-Rad Laboratories Ltd, Watford, UK) in accordance with the manufacturer’s instructions Once conjugated the product was diluted 1 in 10 in HRP Stabilising Buffer (ClinTech, Guildford, UK) and stored un-fractionated in 50μl volumes at −20°C.

### Conjugates for capture and DABA

HRPO-conjugated rZKVNS1Ag was appropriately diluted in conjugate diluent (Initially product kit GE 34/36, (gifted by DiaSorin UK, Dartford, Kent, UK) and then subsequently Clintech Conjugate diluent, (ClinTech, Guildford, UK) and either used as such (unblocked) or diluted in the same conjugate diluent to which had been added a molar excess of rDVNS1Ag of defined serotype (blocked). The performance of the conjugate either blocked or unblocked was compared across the range of dengue sero-positive samples and samples likely to contain anti-ZKV from patients with confirmed ZKV infection.

### Serological controls for capture and DABA

Human IgG1 anti-ZKV virus NS1 supplied by the Native Antigen company (Kidlington, Oxford, UK) was used as a positive control. Used at 10ug/ml in G-capture assay as the positive quality control it was required to give an optical density (OD) between 0.8 and 1.5. Used at 2ug/ml in the DABA assay it was required to give an OD between 2.5 and 3.5.

Pooled normal UK human plasma (NHP) previously screened for blood borne viruses constituted the unreactive control. Test samples reacting greater than the mean negative control +0.1 OD where considered to be reactive for anti-ZKV.

## Results

### Demonstration of cross-reactivity

The antibody reactivity of 147 samples (118 samples from the 2014 outpatient panel and 29 blood donor samples from 2013) archived before the recognised introduction of ZKV into Brazil was determined in the EuroimmunZ indirect Elisa and in the unblocked DABA (Figure 1). Thirty five samples (24%) were reactive on the EuroimmunZ whilst 32 samples (22%) were reactive on the unblocked DABA, 28 (19%) were concordantly reactive in both assays. Fourteen of the 147 (10%) were reactive in the indeterminate zone in EuroimmunZ, one of which was reactive in the unblocked DABA. Ninety eight samples (67%) were un-reactive in the EuroimmunZ, three were reactive in the unblocked DABA.

**Fig 1.**
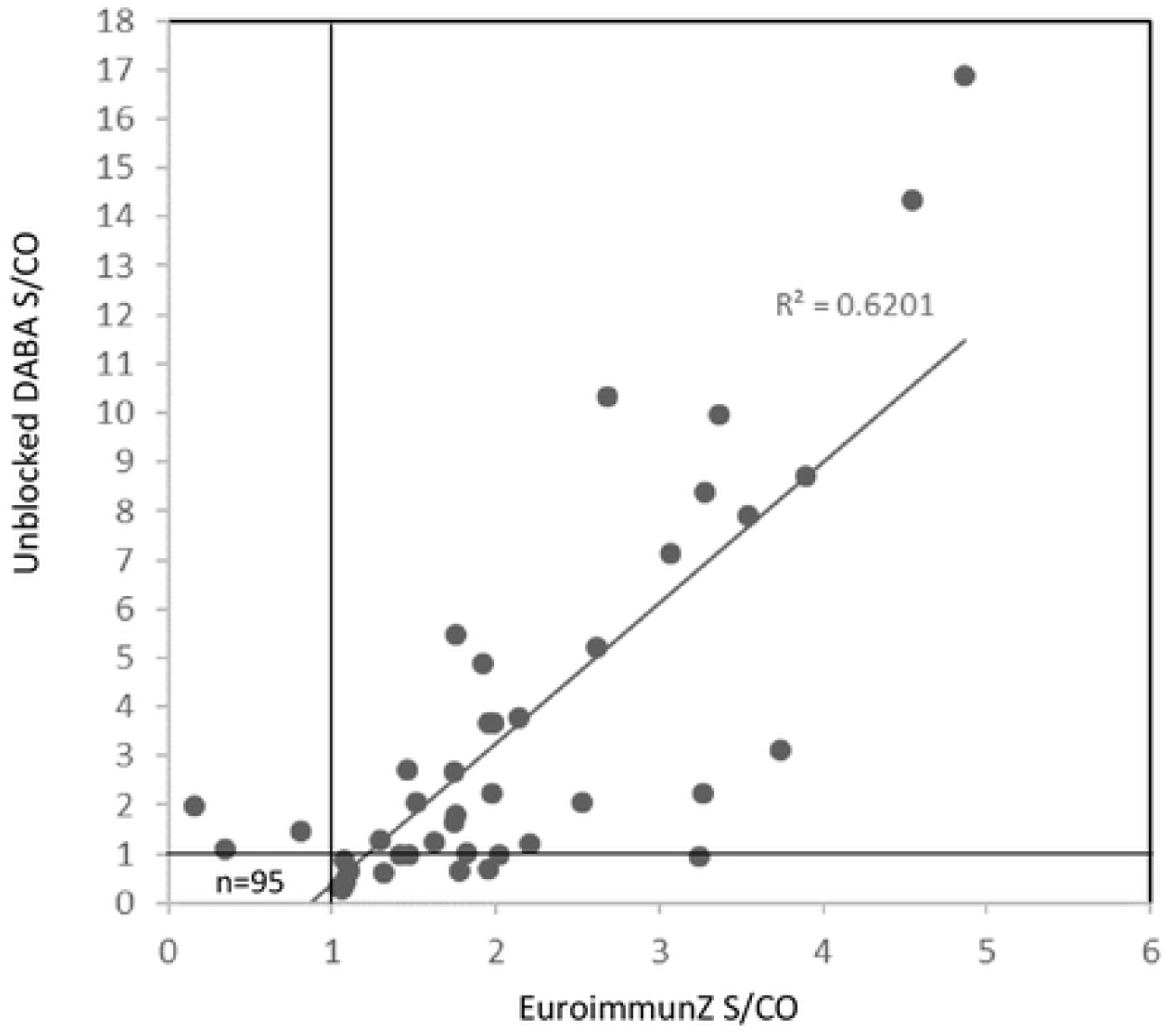
EuroimmunZ vs Unblocked DABA reactivity in pre-Zika era panels. Plot of sample to cut off (S/CO) ratios of 147 pre-Zika era samples tested in un-blocked DABA and EuroimmunZ. Fourteen samples with EuroimmunZ ratios <1.1 were considered indeterminate by EuroimmunZ criteria. Three samples were reactive by DABA alone. Solid lines represent the cut off value for each assay. Trend line is displayed.

The reactivity given by sera containing anti-DV antibody following known infection by Dengue serotypes 1 through 4 was determined in each of the three assays, EuroimmunZ, DABA and IgG capture, both using un-blocked conjugates. The proportion of samples reactive for anti-ZKV in one or more assays ranged from 29 of 32 (91%) of samples from patients with secondary DV2 infections, 9 of 20 (45%) of samples from patients with secondary DV3 to 11 of 40 (28%) of samples from patients undergoing secondary DV4 samples. Seven of 28 (25%) samples from patients with primary DV1 reacted in one or more assays (data not shown).

### Sensitivity of different assay formats for detecting a ZKV serological response

The evolution of detectability of antibody to ZKV in EuroimmunZ, in DABA and in IgG capture using unblocked conjugates, was investigated using a panel of samples from 56 patients undergoing a confirmed acute ZKV infection. Ninety one sera were tested, 69 were reactive in EuroimmunZ, 78 were reactive in DABA and 60 in IgG capture (Figure 2). The majority of the samples (52/91, 57%) were concordantly reactive in all three assays. Ranking the samples by time after the onset of symptoms indicated that of the 22 samples that were taken in the first week of symptoms, seven were reactive in EuroimmunZ, 11 were reactive in DABA and ten in IgG capture. Of the 49 samples taken between two and four weeks after the onset of symptoms, all but one were reactive in both EuroimmunZ and DABA, only 38 were reactive in IgG capture. Of the 20 samples taken more than one year after the acute illness only 15 were reactive in any assay, ten in EuroimmunZ, 14 in the DABA and eight in the IgG capture.

**Fig 2.**
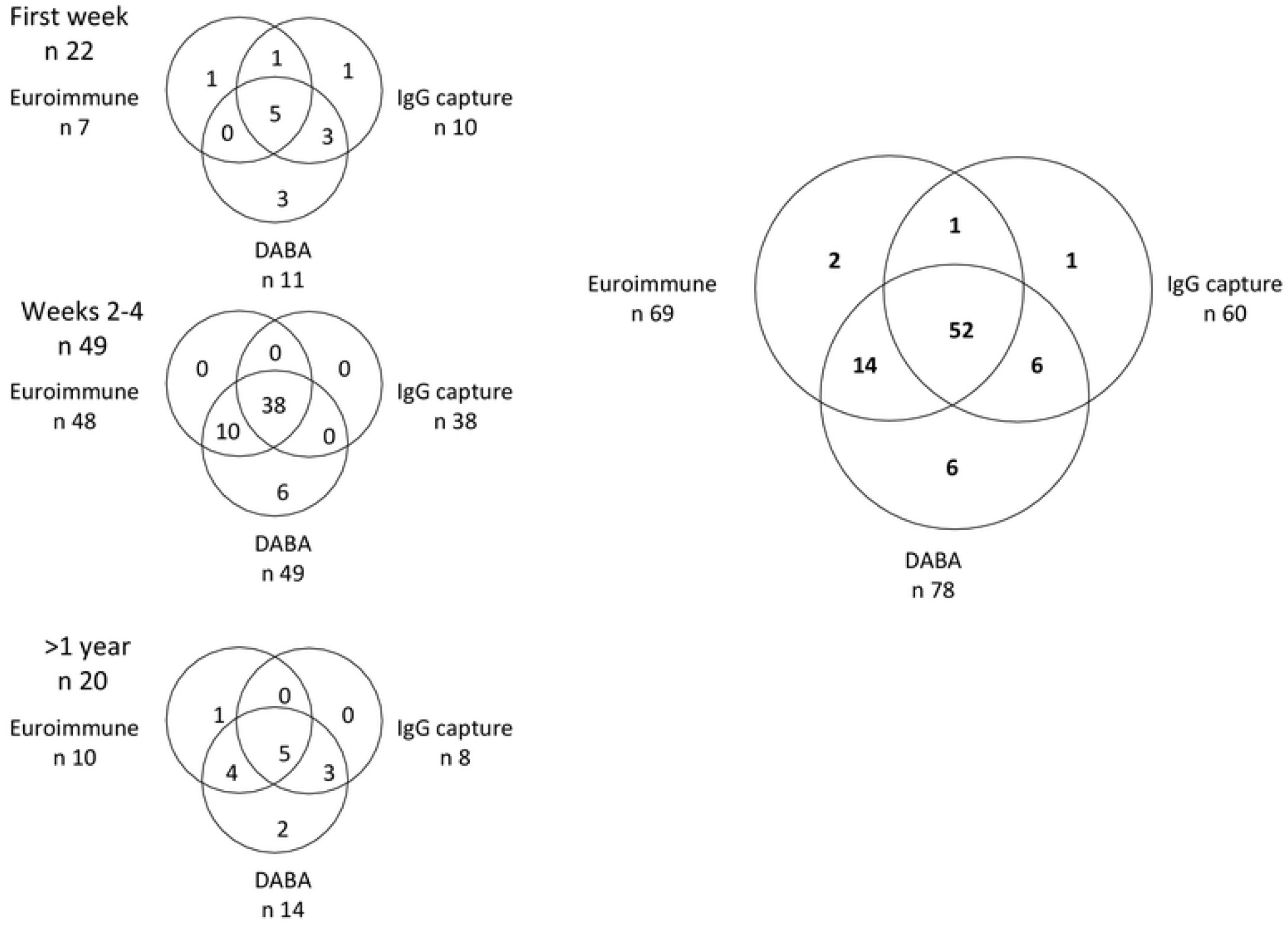
Antibody reactivity across three assays in samples from PCR confirmed Zika cases. Venn plots for 91 samples from patients with PCR confirmed ZKV infection. Samples giving sample to cut off ratios >1.0 were defined as reactive. The right hand plot shows reactivity for the panel of 91 samples tested in the three different assays. The three panels on the left in vertical order show the reactivity broken down into the first week, second to fourth week and one or more year after onset of symptoms. The individual reactivity of samples in each test is shown and the overall reactivity for each assay displayed.

### Conjugate quenching to reduce cross-reactivity

The addition of blocking amounts of unlabelled recombinant DV NS1 antigen was investigated to reduce the signal of apparent Dengue serum antibody reactivity in the two assays, IgG capture and DABA, both using HRPO-conjugated rZKVNS1Ag as a detector. A panel comprising 32 DV antibody-containing sera samples including those that were previously falsely reactive in the unblocked ZKV IgG capture and/or DABA assays was selected. The panel was then retested in both assays using a modified conjugate diluent containing unlabelled rDV3NS1Ag at a concentration of 25μg/ml. A single serum remained reactive in the DABA, all remaining non-specific reactivities were quenched by the use of conjugate diluent containing rDV3NS1Ag (data not shown).

### Sensitivity, specificity and optimisation of blocked conjugates for detecting anti-ZKV

A subset of the panel of 95 sera from ZKV patients were re-tested initially using the rDV3NS1Ag as the sole constituent of a blocked buffer. In the majority of samples, the anti-ZKV reactivity was equivalent in both IgG capture and DABA with both conjugates, however a proportion of samples (six from 59 in DABA and eight from 41 in IgG capture) showed significant reduction of reactivity with the blocked conjugate (circled, figure 3a and 3b respectively).

**Fig 3.**
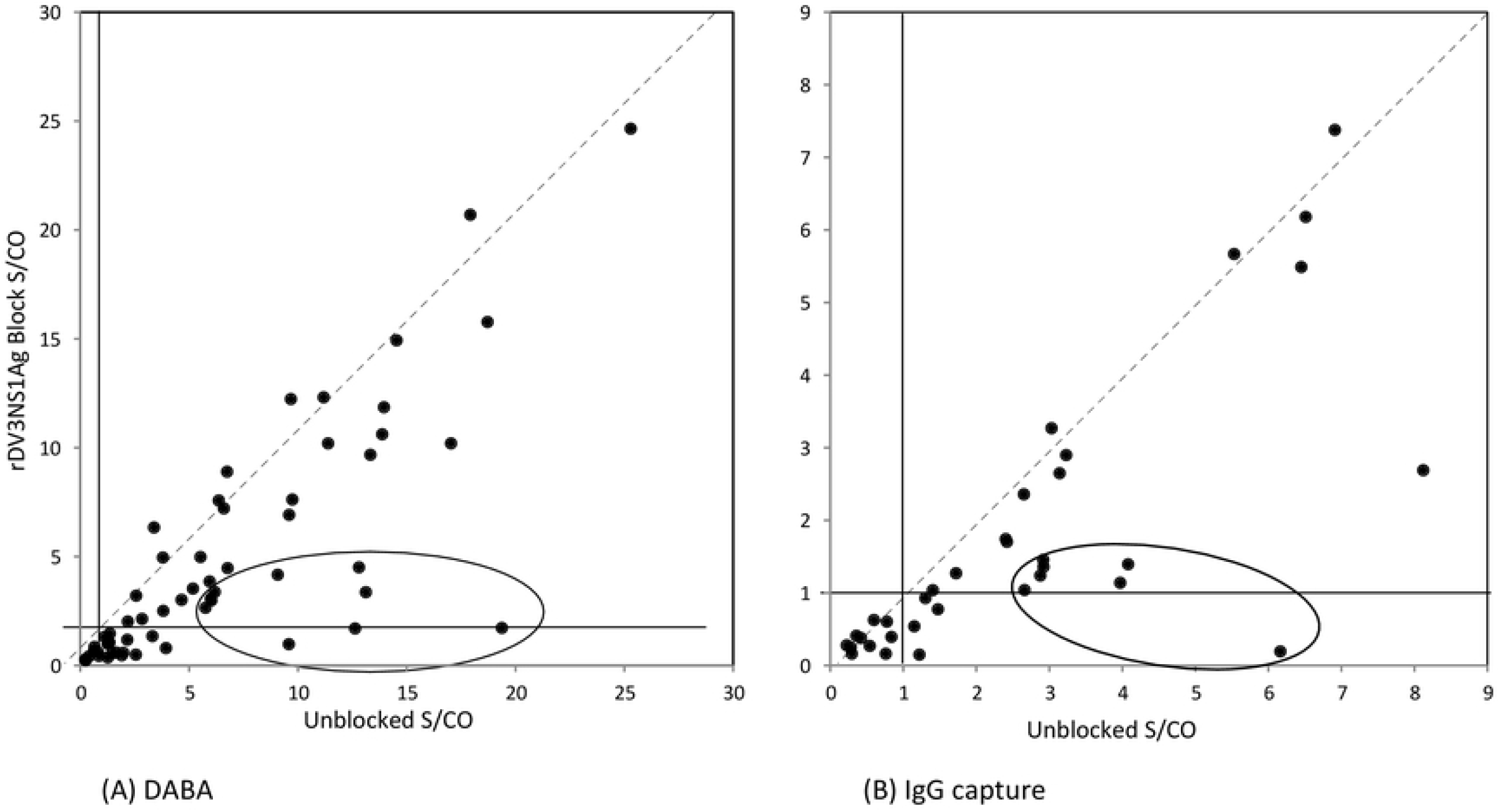
IgG capture and DABA data on Zika confirmed panels comparing DV3NS1Ag quenched and un-quenched conjugates. X by Y plots of sample reactivity from patients known to have been infected with ZKV. Reactivity is expressed as sample to cut off ratios when tested using un-blocked conjugate and rDV3NS1Ag-blocked conjugate. Left hand panel (A) displays the results with 59 ZKV convalescent sera (São Paulo and Rio de Janeiro) tested in the DABA using both conjugates. Dotted line is a line of interpolated equivalence assuming no difference in reactivity. Solid lines represent the assay cut off values. Right hand panel (B) similarly displays the reactivity of 41 samples from patients with confirmed ZKV infection tested in the IgG capture using unblocked and blocked conjugate diluents. Samples from patients with proven ZKV showing a reduced reactivity resulting from the blocked conjugate are circled in both panels.

To investigate in more detail the serotype specificity of the quenching effect of rDVNS1 proteins, a panel of 20 sera previously displaying strong reactivity in the unblocked IgG capture assay was run a second time in the IgG capture and the DABA assay with two different conjugate diluents. A normal conjugate diluent was used for the unblocked version as referent in comparison with the blocked version of the assay using conjugates containing quenching amounts of rDVNS1Ag, present as rNS1Ag of each DV serotype individually. The degree of residual activity displayed by the different quenching rDVNS1Ags was greatly influenced by the rDVNS1Ag serotype (Table 1). We also investigated quenching as a quadrivalent equipotent rNS1Ag-blend of all 4 serotypes at concentrations from 25, 12.5 to 6.25μg/ml. The quadrivalent mix ablated DV-related cross reactivity in the DABA, two samples remained consistently reactive in the IgG capture assay. Specific anti-ZKV control reactivity remained detectable under all quenching mixtures (Table 2). There was some evidence for a titration of the quenching function over the range from 25 to 6.25μg/ml of competitor DVNS1 homologue in the IgG capture conjugate, with mean quenched sample reactivity ranging from 0.63 through 0.74 to 0.87 across the titration of quenching NS1 antigen. A similar trend is seen for DABA with mean reactivities of 0.56, 0.61 and 0.63. A single stochastic low level reaction was seen in the 6.25μg/ml quench DABA assay but the sample was not retested. Based on the relative effectiveness at 6.25μg/ml, this homologue antigen quench concentration was adopted for further studies.

**Table 1.**
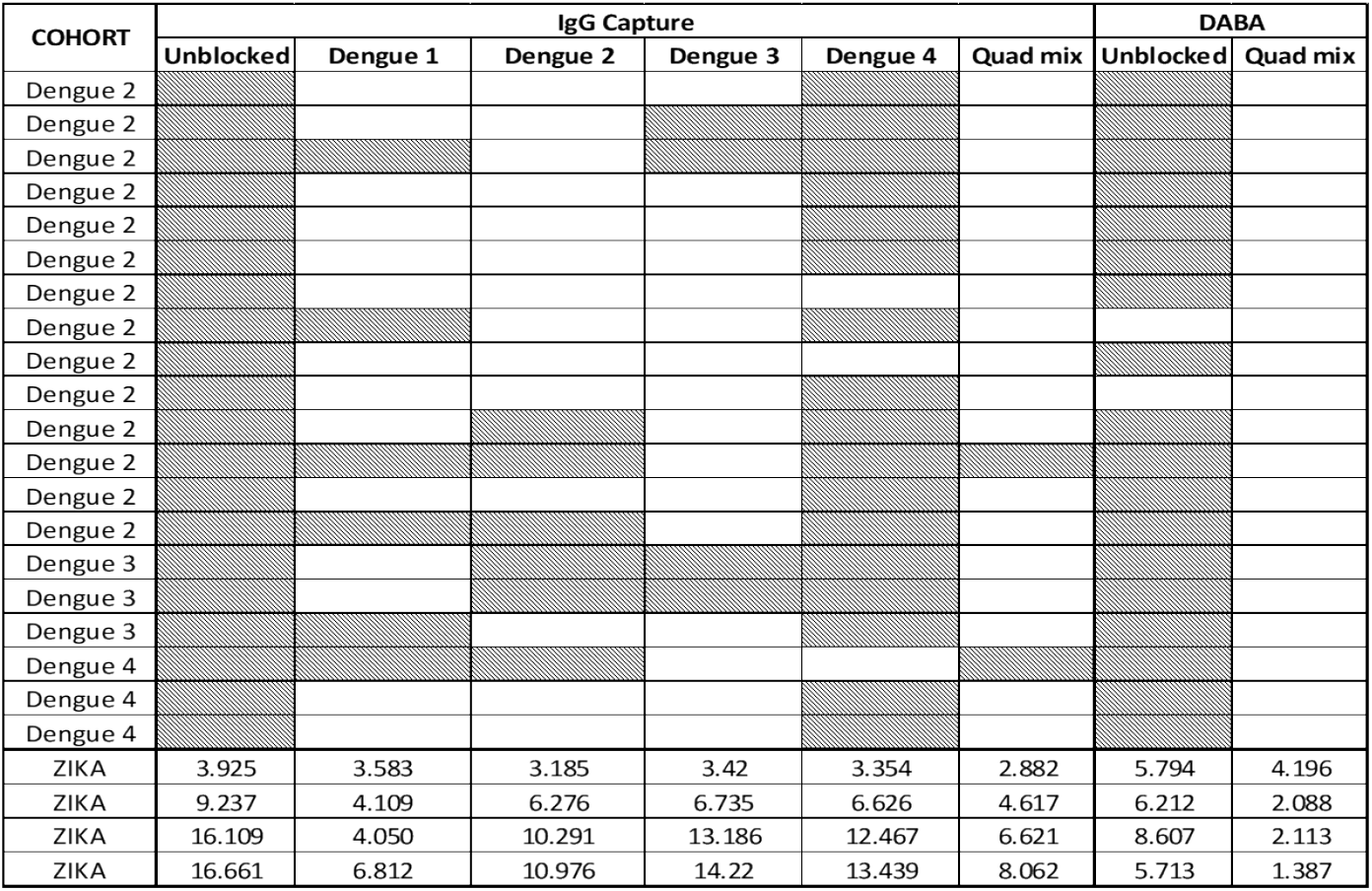
Blocking of non-specific antibody reactivity using a panel of Dengue serotype 1 to 4 antigens. The reactivity of a panel of 20 samples from patients with previous dengue virus infection of known serotype when tested with unblocked and blocked (rDVNS1Ag 12.5μg/ml) conjugate diluents in the IgG capture and the DABA assay. The quenching component in a blocked conjugate is of either a single DV serotype as shown for the IgG capture or of a quadrivalent preparation containing equipotent quantities of all four serotypes as shown for both the IgG and the DABA. Shaded cells represent sample/cut off reactivity >1. Reactivity of the control ZIKA-positive samples was maintained (bottom four rows).

**Table 2.** Titration of un-labelled competitor homologue DV NS1. Optimisation of the quadrivalent un-labelled competitor homologue DV NS1block. Sample /cut off ratios are shown for both the IgG capture assay and the DABA. The sample reactivity unblocked and blocked at the three levels of 25, 12.5 and 6.25μg/ml is shown for each assay, samples still reactive in the face of conjugate blocking are displayed in bold. Reactivity of the control ZIKA-positive samples is displayed (bottom four rows).

| COHORT | IgG Capture |  |  |  | DABA |  |  |  |
| --- | --- | --- | --- | --- | --- | --- | --- | --- |
|  | Unblocked S/CO | Quad Mix 25ug/ml S/CO | Quad Mix 12.5ug/ml S/CO | Quad Mix 6.25ug/ml S/CO | Unblocked S/CO | Quad Mix 25ug/ml S/CO | Quad Mix 12.5ug/ml S/CO | Quad Mix 6.25ug/ml S/CO |
| Dengue 2 | 2.184 | 0.384 | 0.493 | 0.623 | 12.278 | 0.513 | 0.536 | 0.632 |
| Dengue 2 | 8.421 | 0.439 | 0.6 | 0.778 | 6.57 | 0.528 | 0.572 | 0.641 |
| Dengue 2 | 4.379 | 0.544 | 0.754 | **1.472** | 11.858 | 0.491 | 0.557 | 0.632 |
| Dengue 2 | 6.926 | 0.485 | 0.571 | 0.759 | 9.968 | 0.543 | 0.68 | 0.784 |
| Dengue 2 | 8.574 | 0.453 | 0.799 | 0.858 | 6.091 | 0.468 | 0.557 | 0.637 |
| Dengue 2 | 3.011 | 0.475 | 0.526 | 0.594 | 4.608 | 0.445 | 0.546 | 0.582 |
| Dengue 2 | 1.426 | 0.485 | 0.654 | 0.67 | 5.301 | 0.687 | 0.722 | 0.559 |
| Dengue 2 | 4.337 | 0.686 | 0.857 | **1.075** | 0.77 | 0.468 | 0.5 | 0.486 |
| Dengue 2 | 2.279 | 0.457 | 0.447 | 0.467 | 4.421 | 0.347 | 0.433 | 0.426 |
| Dengue 2 | 6.405 | 0.517 | 0.547 | 0.665 | 0.977 | 0.589 | 0.577 | 0.591 |
| Dengue 2 | 4.268 | 0.37 | 0.576 | 0.585 | 8.252 | 0.558 | 0.588 | **1.072** |
| Dengue 2 | 2.642 | **1.573** | **1.723** | **2.024** | 2.214 | 0.408 | 0.536 | 0.431 |
| Dengue 2 | 5.453 | 0.562 | 0.675 | 0.783 | 1.864 | 0.543 | 0.619 | 0.646 |
| Dengue 2 | 6.611 | 0.539 | 0.671 | 0.807 | 9.221 | 0.445 | 0.531 | 0.486 |
| Dengue 3 | 2.021 | 0.59 | 0.621 | 0.693 | 1.95 | **1.245** | 0.938 | 0.948 |
| Dengue 3 | 9.416 | 0.777 | 0.915 | 0.91 | 14.375 | 0.725 | 0.768 | 0.756 |
| Dengue 3 | 3.337 | 0.709 | 0.634 | 0.679 | 2.229 | 0.702 | 0.711 | 0.591 |
| Dengue 4 | 2.226 | **1.207** | **1.379** | **1.429** | 12.66 | 0.506 | 0.588 | 0.49 |
| Dengue 4 | 2.553 | 0.736 | 0.762 | 0.717 | 25.864 | 0.528 | 0.608 | 0.564 |
| Dengue 4 | 9.947 | 0.594 | 0.683 | 0.788 | 1.663 | 0.521 | 0.572 | 0.577 |
| Mean S/CO | 4.821 | 0.629 | 0.744 | 0.869 | 7.157 | 0.563 | 0.607 | 0.627 |
| ZIKA | 3.505 | **3.237** | **2.882** | **3.09** | 5.794 | **5.894** | **4.196** | **3.216** |
| ZIKA | 6.221 | **4.649** | **4.617** | **4.915** | 6.212 | **2.702** | **2.088** | **1.865** |
| ZIKA | 12.621 | **5.888** | **6.621** | **7.887** | 8.607 | **2.694** | **2.113** | **1.746** |
| ZIKA | 12.937 | **7.991** | **8.062** | **9.123** | 5.713 | **4.325** | **1.387** | **2.465** |

To explore further the quenching effect of using conjugate blocked with the optimised quadrivalent mixture of rDVNS1Ag, 87 samples from the 2014 outpatient cohort were retested using quadrivalent blocked conjugate at 6.25ug/ml. Twenty three samples reactive in the unblocked DABA and 27 reactive in EuroimmunZ were all unreactive in the quadrivalent blocked DABA (Figures 4a and 4b). Nine samples giving indeterminate reactivity on the EuroimmunZ were unreactive in the quadrivalent blocked DABA. A further 19 anti-DV reactive samples were subsequently included to give a panel total of 106 samples which were all unreactive in the DABA indicating a high specificity.

**Fig 4.**
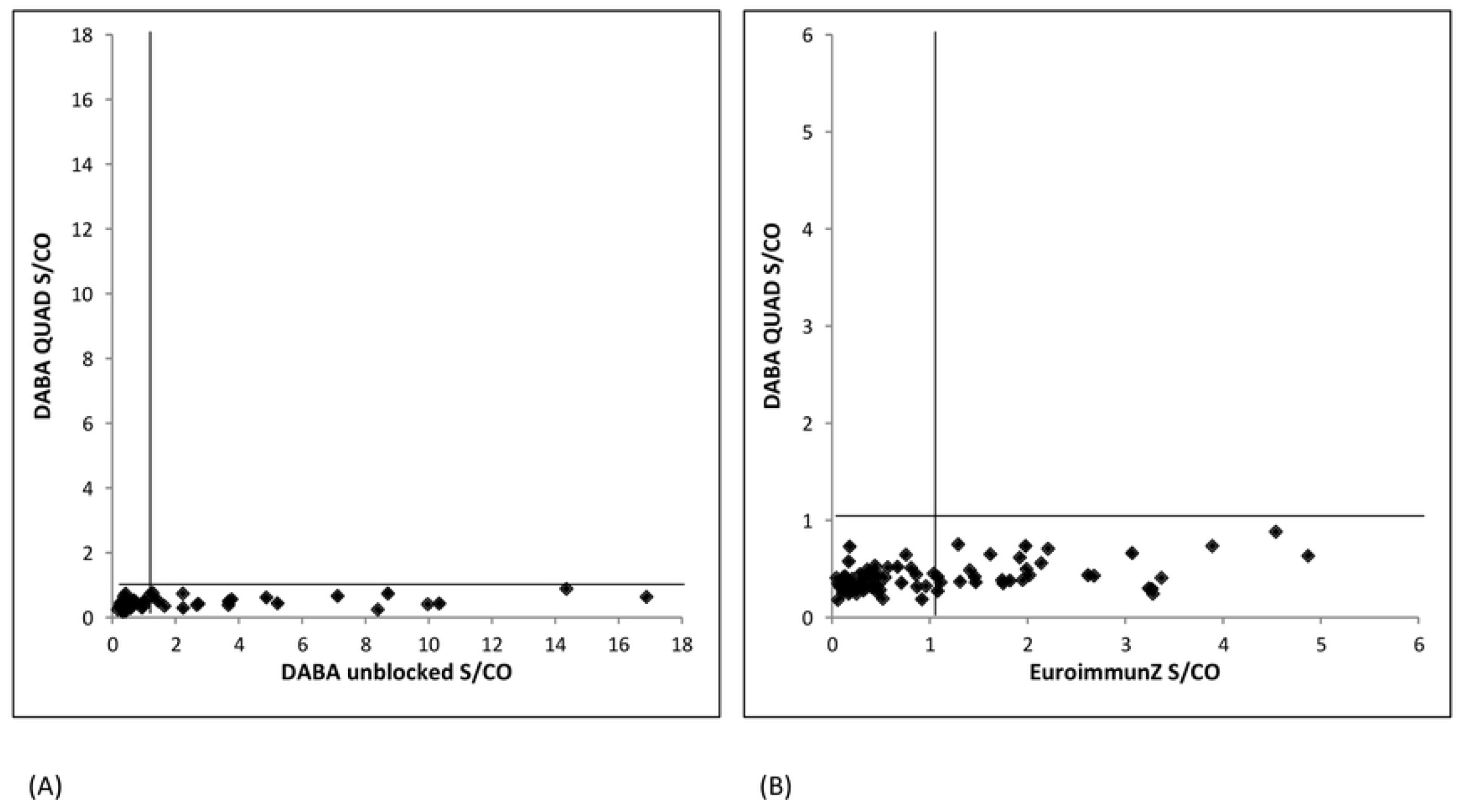
Antibody reactivity in Dengue quadrivalent antigen blocked assay: comparison with unblocked DABA and EuroimmunZ. X by Y plots of reactivity displayed by 87 selected samples (São Paulo and Rio de Janeiro) drawn from the pre ZIKA-era panel expressed as sample to cut off ratios. Left hand panel (A) displays the results with sera tested in DABA using both quad blocked and unblocked conjugates. Right hand panel (B) similarly displays the reactivity of the same samples tested in quad blocked DABA and EuroimmunZ.

A further panel of 38 samples from 19 ZKV-infected patients taken at various times after the onset of disease was again re-tested in the IgG capture assay only (to conserve sample volumes) using the optimised quadrivalent block and the resulting reactivity compared with the reactivity in the EuroimmunZ assay. This demonstrated a considerable divergence between reactivity in the two assays, most prominently observed early in the illness time course with the anti-ZKV reactivity in the IgG capture assay with a blocked conjugate often found to be very much reduced, particularly in the first week after diagnosis (Figure 5).

**Fig 5:**
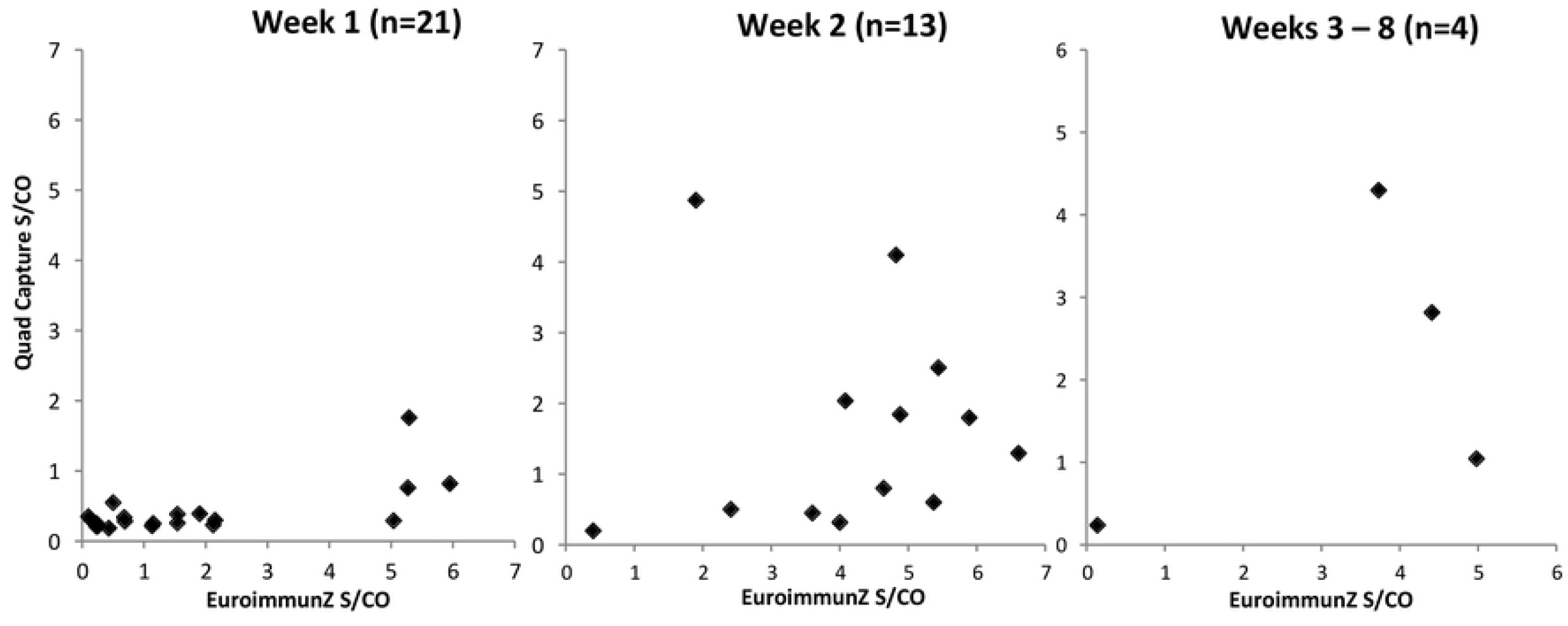
Zika seroconversion panels: EuroimmunZ vs Quadrivalent antigen blocked IgG capture by time from infection. X by Y plots of sample reactivity, expressed as sample to cut off ratios in IgG capture assay using quad blocked conjugate and in EuroimmunZ, of 38 samples grouped by time since onset of symptoms.

## Discussion

After the first description of ZKV virus infection in Africa nearly 80 years ago and the sporadic numbers of human infections identified subsequently associated with mild erythematous illness, the first significant cluster of cases since then was reported from Yap Island in Micronesia in 2007 [2]. By the early part of 2016, the WHO declared that ZKV infection, associated as it was with severe neurological disorders in the new-born, was a public health emergency of international concern. A particular aspect arising from this decision was the requirement for specific and sensitive serological assays for the detection of anti-ZKV antibody. This was always going to be challenging because of the potential exposure to multiple co-circulating Flavi-viruses in endemic areas, with the known potential to elicit serological cross reactivity between these infections. Particular assay formats have individual attributes, an observation which underlines the value of taking a broader view when setting out to develop serological tests for infectious agents. This approach is exemplified by our recent experience with Ebola virus serology [13] and echoed by Balmaseda and colleagues [12]. Recognising that antibody to envelope proteins is often cross-reactive within subgroups of the Flavi-viridae family, Stettler and her colleagues [14] moved to use the non-structural protein NS1 in view of data suggesting that this protein was likely to provide a more species-specific antigen moiety for use as the solid phase in immunoassays. A similar conceptual approach is employed in the benchmark and widely used EuroimmunZ assay which uses rZKVNS1Ag in an indirect assay format. These observations informed our choice of the NS1 target and our decision to explore assay formats other than the indirect immunoassay.

It is clear however that the use of the NS1 protein in the EuroimmunZ indirect immunoassay format is still associated with significant cross-reactivity likely to be generated by prior Flavi-virus co-circulation and repeated exposure in the community prior to the introduction of ZKV into new geographical areas (Figure 1). In examining the different assay formats of IgG capture and DABA we find a similar lack of high specificity from patient and blood donor samples collected in two different areas of South East Brazil before the introduction of ZKV into the Americas.

The comparative sensitivity of the three assay formats was examined using 91 samples from patients sampled at different times after the onset of disease symptoms (figure 2) using unblocked conjugates in the IgG capture and DABA formats. In overall ranking it is likely that the DABA format, in essence a total antibody test, was marginally more proficient at detecting early and late serological markers than the other two formats. The DABA format will recognise antibody of any class and does indicate the potential of being a total antibody assay of high sensitivity. It is however questionable exactly what reactivity was being detected, as false reactivity was also displayed by all three formats in samples from persons who had previous DV infection. This was most notable after secondary DV2 infection and all three test formats were variously susceptible including those using a labelled antigen conjugate. We therefore investigated a novel approach of including unlabelled Dengue virus antigens in the conjugate diluent. In the first instance one antigen serotype rDV3NS1Ag at 25μg/ml was used in a blocked conjugate. This was effective in quenching nonspecific reactivity in both formats but reduced the reactivity of ZKV convalescent samples (Figure 3). In many cases there was concordant reduction in both assay formats.

Our data showed that extinguishing cross reactivity was best achieved by using an equipotent four component mix of rDVNS1Ags (Table 1). Optimisation indicated that and this was used in all subsequent analyses at 6.25μg/ml. It was effective in quenching false reactivity of the pre-ZKV <u>period</u> samples (Figure 4). Using a panel of 35 samples from patients taken at various times following confirmed ZKV diagnosis, a significant and notable reduction in the magnitude of the ZKV-specific signal was seen again, particularly early in the course of the antibody response (Figure 5). This indicated that a considerable component of the early antibody reactivity, irrespective of assay format, may be related to prior cross reacting dengue or other related flavi-virus antibody. This is not altogether surprising since not only does ZKV infection boost dengue antibody [15] but the concept of “original sin” can also be applied to flavi-virus reinfection [16]. Taken together these two data sets generated by using blocked conjugates suggest that ZKV infection may induce an anamnestic boost to DV antibody from previous Dengue virus exposure.

Such observations again underline the need for a serological strategy to minimise cross reactivity and reveal ZKV-specific responses which can easily be masked by cross reaction [17]. Approaches to absorb out interfering reactivity prior to analysis provide one option, so too does the use of residue deletion to remove cross reacting epitopes in recombinant proteins. It is likely that the attribute of using assay formats that allow quenching of the conjugate as described here will deliver specificity whilst retaining sensitivity. It also allows the inherent sensitivity of the DABA format to be used without the usual trade off between sensitivity and specificity. The additional advantage of this for reverse immunoglobulin capture assays is considerable as it opens up not only a potential increase in specificity for IgM detection but also the application of serology to non-blood analytes especially oral fluid, a long established methodology [18], which remains of considerable value in field epidemiology as we have found with other emerging infections including Ebola [19].

In summary there remains a clear need for better ZKV serological tools for diagnostic and epidemiological purposes; the data presented here support the utility of the assay formats developed and it is now important to generate more comprehensive field data in order fully to validate the application of the DABA and the Capture assay as described and provide more substantive sensitivity and specificity data in the first instance. Whilst active transmission has declined substantially across the Americas, much remains to be elucidated about the natural history of ZKV virus infection. Cases are being sporadically described elsewhere, with potential for rapid spread where environmental and population conditions are favourable. Efforts to develop a ZKV vaccine also continue. In this context, affordable, highly specific and sensitive antibody tests are needed to expand the testing capacity in areas that have been or may become affected by ZKV virus.

## Acknowledgements

We would like to thank colleagues in Diasorin for the kind gift of the conjugate diluent. DB is partially funded by European Union’s horizon 2020 research and innovation programme (grant agreement no. 734857), Zikaction and Zikaplan consortia (grant agreement 734584). RMC is affiliated to the NIHR HPRU in Emerging and Zoonotic Infections at University of Liverpool in partnership with PHE, in collaboration with LSHTM

